# Changes in the urine proteome after massage in healthy people

**DOI:** 10.1101/2023.02.23.529641

**Authors:** Bao Yijin, Pan Xuanzhen, Gao Youhe

## Abstract

This study aimed to explore the effect of massage on the urine proteome of healthy people. In this study, participants underwent 1-hour whole body massage. Urine samples were collected at 0, 2, and 24h after the massage and urine proteins were analyzed using liquid chromatography-tandem mass spectrometry (LC-MS/MS). Compared with the control (before massage), 41 differential proteins were identified in the group 2h after the massage, the random mean number of differentially produced proteins was 11 with 73% confidence, and the biological process of protein enrichment was catecholamine biosynthesis, which was related to the promotion of metabolism and the regulation of neural activities. While 29 differential proteins were identified in the group 24h after the massage, the random average number of differential proteins produced was 10, with the confidence of the difference decreased to 65%, and the effective biological process could not be enriched at this time. The results suggested that the difference in urine protein was greater at 2h after the massage and gradually decreased at 24h after the massage. The proteome of urine may reflect changes in the body following minor massage stimuli, providing a potential way to evaluate the effects of massage therapy.

## 1. Introduction

Massage is a kind of physical medical means, which uses various techniques such as pressing, rubbing, pushing, holding, rolling, and kneading, to promote blood circulation and dredge the channel. In recent years, the Ottawa Experts Panel has proposed another definition of massage, “manipulation of soft tissues and joints by hand or handheld device” ^[1]^. This definition also includes the use of mechanical equipment and emphasizes the reduction of physical pain through massage techniques such as the use of hands or various instruments. Through mechanical stimulation, massage has a direct deformation effect on the tissues and cells of the operation site, activates cell signaling pathways, and produces biological and chemical effects.

Several studies have shown that massage has a positive effect on a variety of diseases, including depressive diseases such as depression and autism, pain syndromes such as arthritis and fibromyalgia, and autoimmune diseases such as asthma and multiple sclerosis ^[1,2]^. However, the specific mechanism of this positive impact is unclear. The effects of massage on the body are multifaceted and complex, and pressing on soft tissues such as muscles not only relaxes the muscles but also increases the neural activity at the level of the spinal cord and the subcortical nucleus ^[3]^, resulting in pain and pleasure ^[4]^, which necessarily involves the activation of multiple pathways. It is known that massage can cause a variety of physiological responses, including accelerated lactic acid clearance, increased lymphatic flow, activation of pathways to promote mitochondrial production, decreased cortisol, and increased serotonin and dopamine ^[1,5]^, which may be the reasons why massage relieves fatigue and makes people feel happy. In the past, most of the evaluations of the massage effect relied on physiological and biochemical indicators or scale evaluation. This study evaluates the effect of massage on the body more comprehensively and objectively through proteomics, detects subtle changes in biological processes through protein expression and content change analysis, and further elucidates the mechanism of action of massage. As a non-invasive and easily available biological fluid, urine is gradually becoming an ideal sample for proteomics research. Urine is not regulated by homeostatic mechanisms and can accommodate and accumulate more changes in the body ^[6]^. In addition, the complexity of the urine proteome is relatively low, and it is easier to detect the changing characteristics of low-abundance proteins ^[7]^, which have the potential to respond sensitively to changes in the body. Therefore, the urine proteome can more sensitively reflect the influence of the whole body by massage, especially small stimuli.

This study described for the first time the changes in urine proteomics before and after massage in healthy people, collected urine before and after massage for non-labeled quantitative proteomic analysis, explored the impact of massage on the body, and provided new clues for the study of the mechanism of action of massage.

## 2. Materials and methods

### 2.1 Collection of urine samples

Six volunteers (6 females, 22-26 years old) received massages at a nearby massage parlor, and a masseur was selected to perform a full-body massage for one hour at moderate strength that the volunteers could bear. Volunteers’ urine was collected before the massage, followed by urine 2 h and 24 h after the massage. The volunteers were adolescents who were healthy and did not suffer from any diseases. The study was reviewed and approved by the ethics committee of the School of Life Sciences of Beijing Normal University, and the volunteers signed the informed consent form.

In this study, the urine samples retained before the massage were the control group, and the urine samples retained after 2h and 24h of massage were the experimental group, and the time was recorded as T1 and T2 respectively. A total of 18 samples were taken. Urine was centrifuged at 3,000×g for 30 minutes and stored in a −80 °C freezer.

### 2.2 Urine protein extraction and proteolytic hydrolysis

A urine sample of 4 ml was removed, thawed, and centrifuged at 4°C, 12,000×g for 30 min. Cell debris was removed, and the supernatant was taken, precipitated with 3 volumes of ethanol overnight, and then centrifuged at 12,000×g for 30 min. The protein precipitation was resuspended in the lysate (8 mol/L urea, 2 mol/L thiourea, 25 mmol/L dithiothreitol, and 50 mmol/L Tris). Protein concentration was measured using the Bradford method. Urine proteolysis was performed by using filter-assisted sample preparation (FASP) methods ^[8]^. Urine protein was loaded onto the membrane of a 10kDa ultrafiltration tube (PALL Corporation) and washed twice with UA (8mol/L urea), 0.1mol/L Tris-HCl (pH 8.5) and 25mmol/LNH_4_HCO_3_ solutions, then 20 mmol/L dithiothreitol (Dithiothreitol, DTT, Sigma) was added for denaturation at 37°C for 1 h, and alkylated with 50 mmol/L iodoacetamide (IAA, Sigma) in the dark for 30 min, with UA and NH_4_HCO_3_ solution washing twice. After that pancreatic enzyme was added at a ratio of 1:50 (Trypsin Gold, Promega, Fitchburg, WI, USA), and incubated overnight at 37°C. After centrifugation overnight, the filtrate after enzymatic hydrolysis was collected as a peptide mixture. The peptides were desalted by HLB column (Waters, Milford, MA) and drained with a vacuum dryer and stored at −80 °C.

### 2.3 LC-MS/MS tandem mass spectrometry analysis

The peptide was redissolved with 0.1% formic acid water, the peptide concentration was determined using the BCA kit, and the peptide concentration was diluted to 0.5 μg/μL. Mixed peptide samples were prepared by 9ul per sample and separated using the High pH Reversed-Phase Peptide Isolation Kit (Thermo Fisher Scientific) as directed. Ten parts of the Fractions were collected by centrifugation, drained using a vacuum dryer and reconstituted with 0.1% formic acid water. iRT (Biognosis Corporation) was added with a 10:1 volume ratio of sample: iRT. Each sample (single experimental sample and ten Fractions) was taken 1ug using the EASY-nLC1200 chromatography system (ThermoFisher Scientific, USA) for data acquisition. The parameters were set as follows: the elution time was 90 minutes, and the elution gradient was mobile phase A: 0.1% formic acid; mobile phase B: 80% acetonitrile. The eluted peptides were detected by the Orbitrap Fusion Lumos Tribird mass spectrometer (Thermo Fisher Scientific, USA). Data dependent (DIA) mass spectrometry data acquisition was performed on all samples, with each sample collected in 3 replicates.

### 2.4 Data analysis

The 10 components separated from the reversed-phase column were acquired in DDA mode and the results were imported into the Proteome Discoverer software (version 2.1) for search. The PD search results were used to establish the DIA acquisition method, calculating the window width and number based on the m/z distribution density. Individual peptide samples were acquired in DIA mode for mass spectrometry data. Mass spectrometry data were processed and analyzed using Spectronaut X software. The raw files collected by each sample DIA was imported for searching. The q value of a peptide less than 0.01 was considered a high confidence protein standard, and protein quantification was performed using the peak area of all fragments in the second peptide.

### 2.5 Statistical analysis

Each sample was subjected to 3 technical replicates and the obtained data were used for statistical analysis. Urine proteins identified before and after gavage were compared for differential protein screening. The conditions for screening differential proteins were as follows: Fold change ≥2 or ≤0.5, and the *P* value of the two-tailed paired t-test <0.01.

### 2.6 Random Grouping Analysis

A total of 12 samples in the T1 group after massage (n=6) and T0 group (n=6) before massage were randomly divided into two groups, and the average number of differential proteins for all random times was calculated according to the same screening conditions in a total of 462 random combinations. The operation of T2 group (n=6) after the massage and T0 group (n=6) before the massage was the same as above.

### 2.7 Bioinformatics analysis

Unsupervised clustering analysis (HCA) was performed using the Wukong platform (https://www.omicsolution.org/wkomic/main/) ^[9]^, and DAVID 6.8 (https://david.ncifcrf.gov/) was used to perform functional enrichment analysis of the identified differential proteins in three aspects: biological processes, cell localization, and molecular function^[10]^. The function of differential proteins was searched in reported studies based on open databases (https://pubmed.ncbi.nlm.nih.gov).

## 3 Results and analysis

### 3.1 Changes of urinary proteome before and after massage

After the end of the test, a total of 18 urine protein samples collected (before and after massage) were analyzed by LC-MS/MS tandem mass spectrometry. A total of 3,917 proteins (2 specific peptides≥ with protein level FDR <1%) were identified and analyzed for unsupervised clustering, from which samples before and 2 h after massage could be roughly distinguished, but samples before and 24 h after massage could not be distinguished. Figure 1 shows the results of a specific sample of unsupervised clustering.

**Figure 1.**
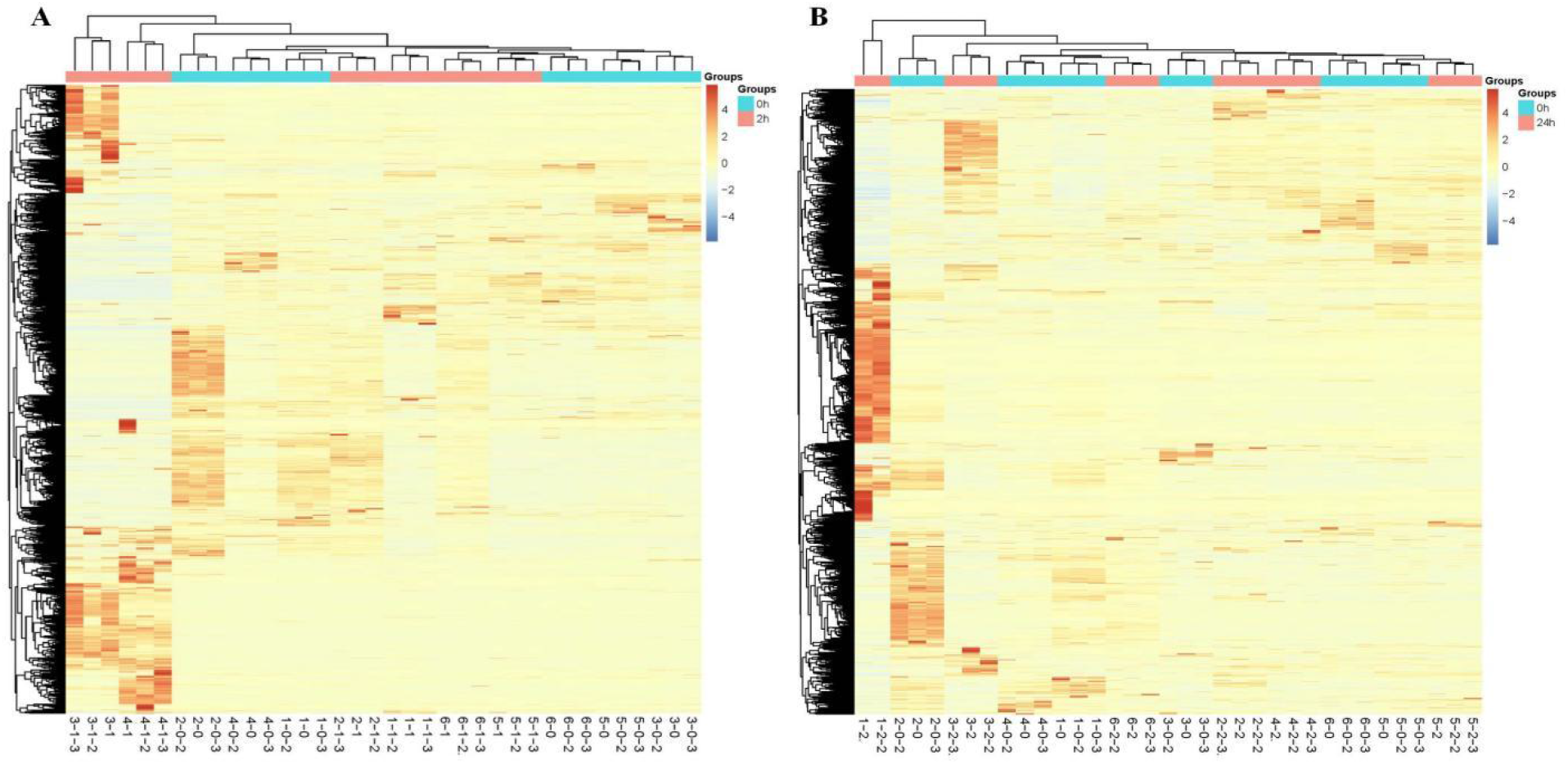
Unsupervised cluster analysis of all proteins in control and experimental groups. (A) 0-2h. (B)0-24h.

The urine protein of T1 and T2 was compared with T0 by self-control method, and the screening criteria were as follows: intergroup variation factor ≥ 2 or ≤0.5, *P*<0.01, the average number of protein spectra in the high abundance group was ≥3, and the protein spectrum of each sample in the high abundance group was higher than that of the low abundance group. Relative to the T0 moment, 41 differential proteins were identified at the T1 moment, of which 3 were upregulated and 38 were down-regulated. Twenty-nine differential proteins were identified at the T2 moment, with 1 up-regulated and 28 down-regulated. More differential proteins were identified at T1 than at T2. Details of the differential proteins identified at different time points are listed in Table 1 (T1 moment) and Table 2 (T2 moment).

**Table 1.**
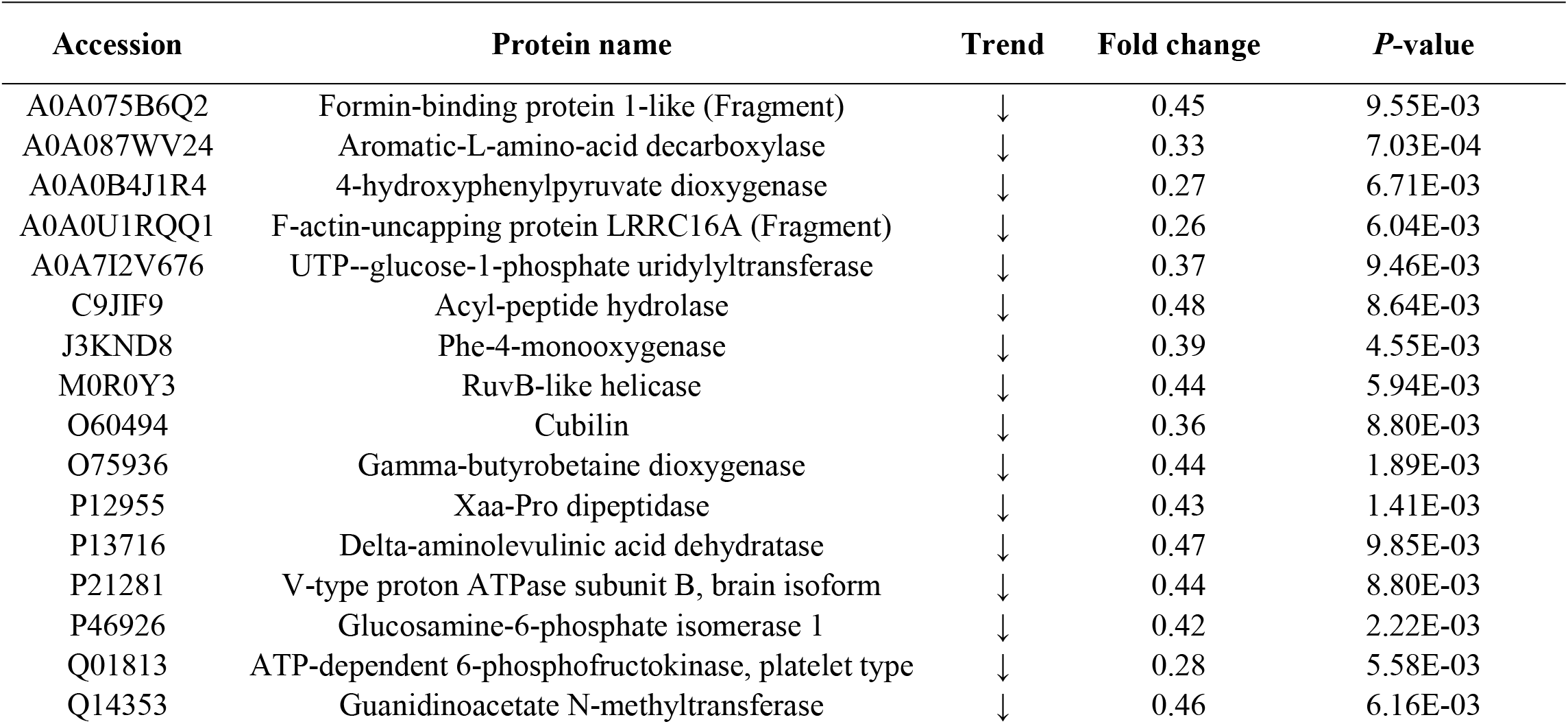
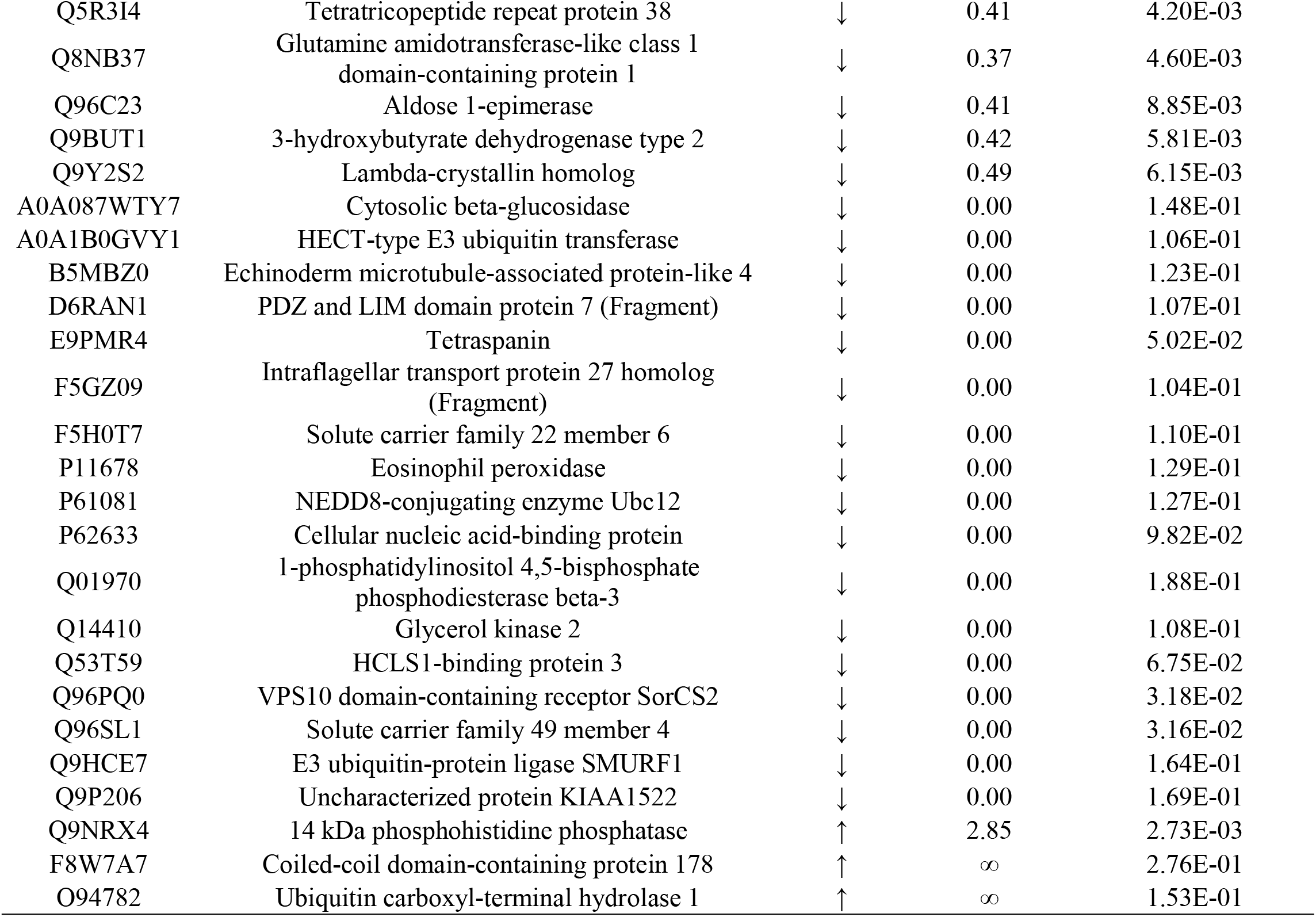
The differential proteins identified at T1.

**Table 2.**
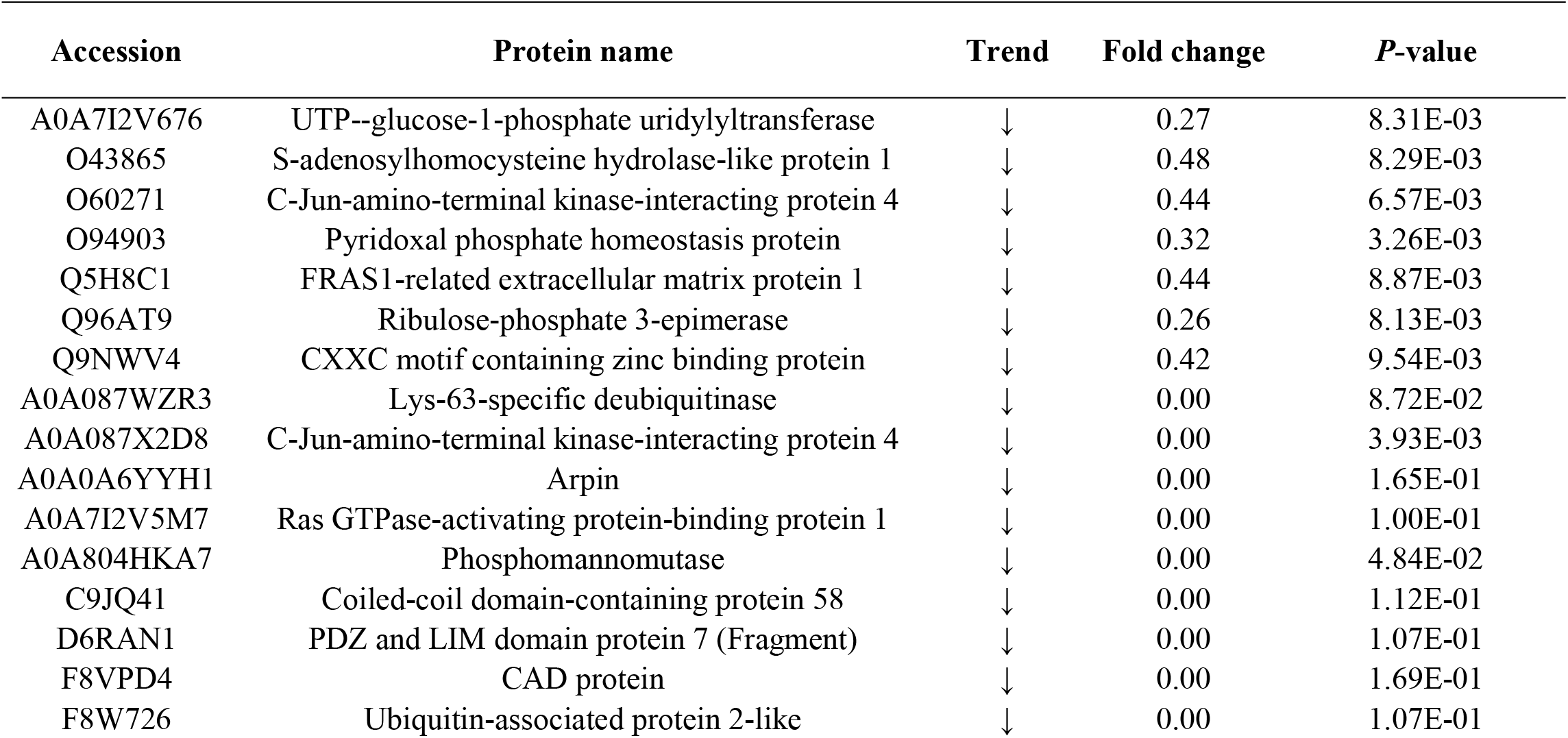
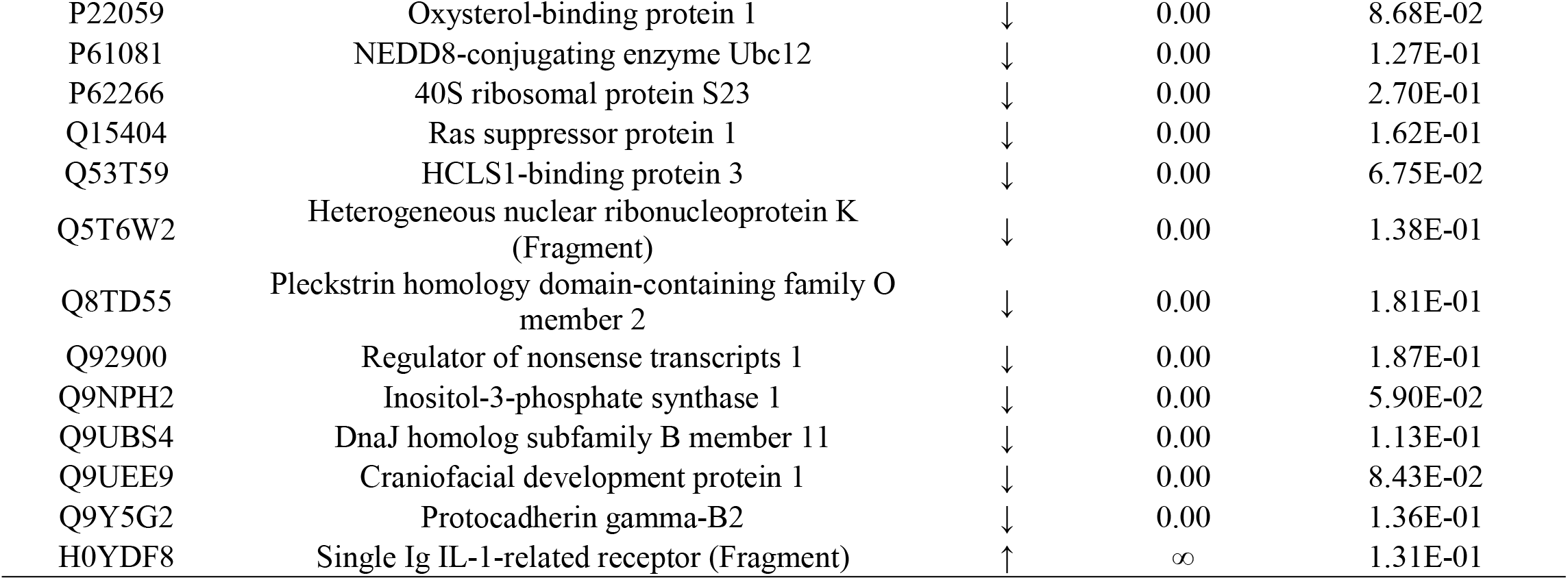
The differential proteins identified at T2.

### 3.2 Results of randomization of urine samples

Given that the number of proteomic features identified in the sample was higher than the sample size, the differences between the two groups might be random. A randomization statistical analysis strategy was developed to confirm whether these differential proteins were induced by massage. A total of 12 samples in the massage T1 group (n=6) and the T0 group before massage (n=6) were randomly divided into two groups, and in a total of 462 random combinations, the average number of differential proteins for all randomized times was calculated according to the same screening conditions as 11 (see Appendix Table 1 for details). These results suggested that only 11 differential proteins could be randomly generated, further indicating that 73% of differential proteins are reliable. The T2 group after massage (n=6) and T0 group before massage (n=6) were calculated according to the above operations, and the average number of differential proteins for all random times was 10 (see Table 2 for details), which showed that 65% of differential proteins were reliable, indicating that the reliability of differential proteins in T2 group after the massage was low.

### 3.3 Functional annotation of different proteins between groups

Based on the above analysis, it is known that proteins 2h after massage showed a more significant difference than those before the massage. Therefore, the DAVID database (https://david.ncifcrf.gov/) was further used to perform functional enrichment analysis of the differential protein identified at the T1 moment in terms of biological process, cellular composition and molecular function (Figure 2). As can be seen from the figure, these differential proteins are mainly involved in the catecholamine biosynthesis process. In terms of cellular composition, most of these differential proteins came from extracellular and cytoplasm, and in molecular functions, we found that most of these differential proteins have functions such as protein binding, oxidoreductase activity, and ubiquitin-protein transferase activity. To identify the major metabolic pathways involved in differential proteins, KEGG pathway enrichment analysis was performed (Figure 2). The results showed that a total of 3 metabolic pathways were significantly enriched, including metabolic pathway, phenylalanine metabolism, and galactose metabolism.

**Figure 2.**
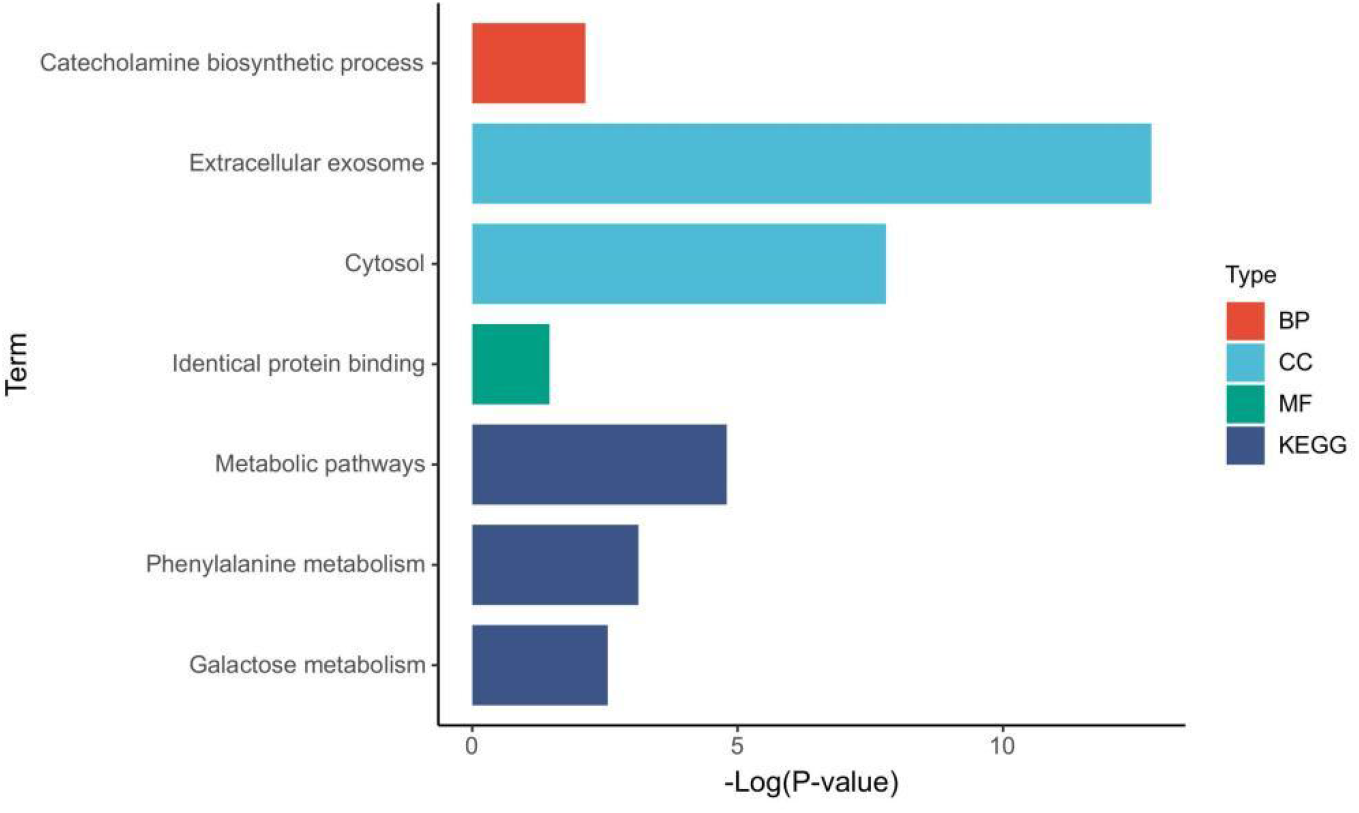
Functional enrichment analysis of differential proteins at T1. (A) Biological process. (B) Cellular component. (C) Molecular function. (D) Pathways.

This showed that massage caused a series of reactions in the body, which could affect the changes in the urine proteome after 2 hours, and with the disappearance of massage stimulation after 24 hours, the changes in the urine proteome also disappeared.

### 3.4 Changes of urine proteome before and after massage in individuals

To explore whether the urine proteome changes before and after massage were consistent, the urine protein identified at the T1 moment and the T0 moment of the individual was compared. The screening conditions for differential proteins were as followed: intergroup variation factor ≥2 or ≤0.5, and *P*<0.01 of the two-tailed paired t-test. The screening results of differential proteins were as follows: a total of 851 differential proteins were screened in No. 1, of which 294 proteins showed an up-regulated trend and 557 proteins showed a down-regulated trend; a total of 383 differential proteins were screened in No. 2, of which 26 proteins showed an up-regulation trend and 357 proteins showed a down-regulated trend; a total of 428 differential proteins were screened in No. 3, of which 168 proteins showed an up-regulation trend and 260 proteins showed a down-regulation trend; a total of 694 differential proteins were screened on No. 4, of which 261 proteins showed an up-regulated trend and 433 proteins showed a down-regulated trend; a total of 383 differential proteins were screened on No. 5, of which 74 proteins showed an up-regulated trend and 209 proteins showed a down-regulated trend; and a total of 443 differential proteins were screened on No. 6, of which 340 proteins showed an up-regulated trend and 103 proteins showed a down-regulated trend.

### 3.5 Functional annotation of individual differential proteins

Further analysis of the biological processes involved in the differential proteins identified at the T1 moment of the individual (Figure 3) showed that proteolysis was enriched in 6 individuals, and 3 biological processes were enriched in 5 individuals, namely glycolysis, reaction to hydrogen peroxide, and carbohydrate metabolism. Therefore, the biological process involved in the differential protein 2h after the massage of each individual was mainly manifested as the metabolism of sugar and protein.

**Figure 3.**
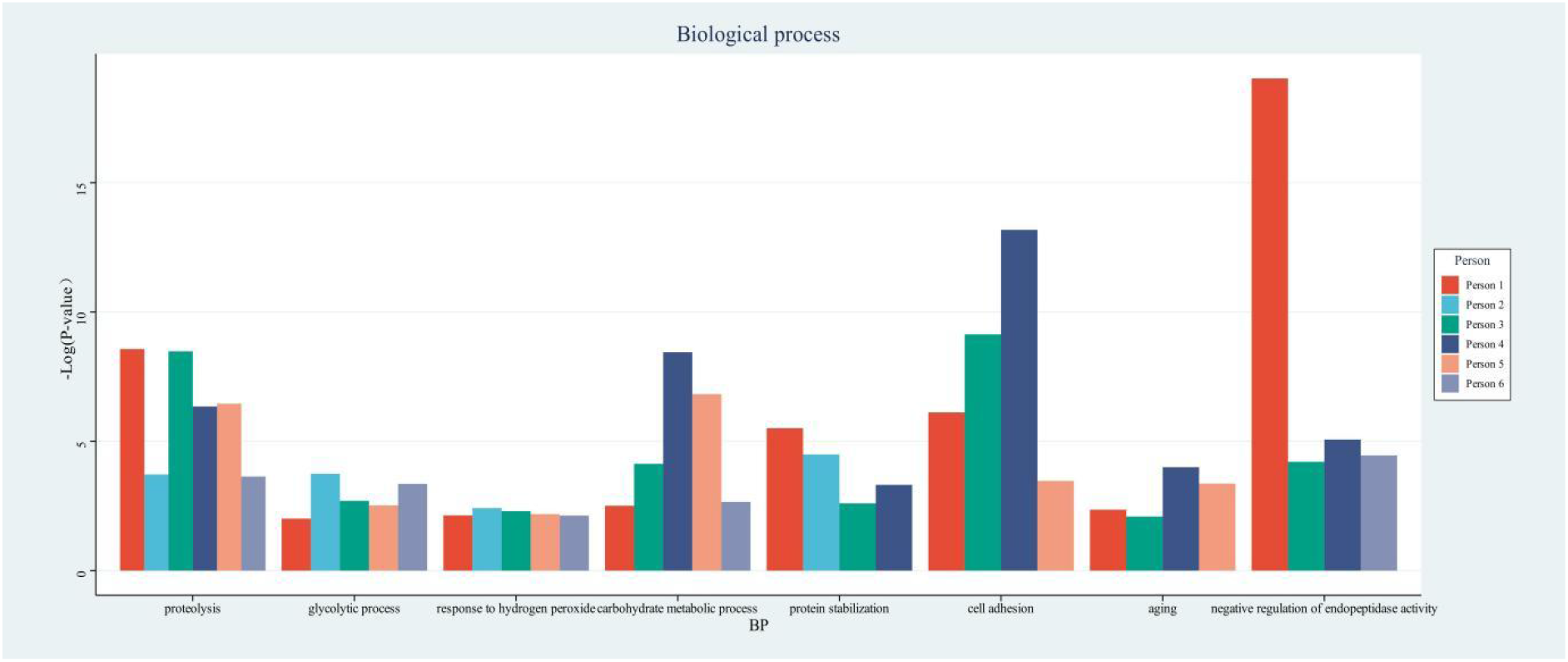
Biological process analysis of differential proteins at T1.

## 4. Discussion

The biological process of enrichment of the T1 moment differential protein after the massage is the catecholamine biosynthesis process, while the T2 time differential protein is not enriched into any biological process. Among them, catecholamines (CAs) are a series of neurological substances containing catechols and amines ^[11]^, mainly including epinephrine (E), norepinephrine (NE) and dopamine (DA). Several studies have found that massage can cause decreased blood levels of corticosteroids ^[12]^, decreased levels of norepinephrine ^[13]^, increased concentrations of serotonin and dopamine ^[2]^, reduced anxiety and improved mood in the massaged subjects. This shows that massage stimulation in this study does cause a series of reactions in the body, promote metabolism, accelerate circulation in the body, and lead to changes in the urine proteome. The period after the massage is the body’s “post-injury repair” process, during which the internal circulation remains in a state of accelerated promotion of repair, and the body can be restored after a period of time. Meanwhile, studies^[14,15]^ have confirmed that massage therapy can play a good role in alleviating and eliminating exercise fatigue, effectively improving exercise fatigue indicators, promoting the recovery of exercise fatigue, preventing delayed muscle soreness, and the immediate effect is obvious. This suggests that the urine proteome has the potential to be used by massage studies related to eliminating exercise fatigue and promoting the mechanism of action of exercise recovery.

In addition, this study also shows that urine sensitivity is far beyond our previous understanding, even if the body is short-lived, micro-stimuli, urine proteome can quickly show changes and enrich into related biological processes, but this change will gradually disappear with the disappearance of small stimuli. Therefore, when carrying out research involving urine sensitivity, it is necessary to collect urine as early as possible. The urine protein group does not change after stimulation, and the urine collection time is likely late. Similarly, the results of this experiment increase the confidence that urine can reflect small changes in the early stages of the disease, which is a strong argument that urine can be used as a source of a new generation of biomarkers.

## 5. Conclusions

This study for the first time described the effects of massage on the urine proteome of healthy people and proved that massage caused a series of reactions in the body through mechanical stimulation, including sugar and protein metabolism, neurotransmitter secretion, and neural activity regulation. These results provide new ideas for the study of the mechanism of action of massage therapy to relieve exercise fatigue and promote sports injury repair.

